# Automated Segmentation of Amyloid-*β* Stained Whole Slide Images of Brain Tissue

**DOI:** 10.1101/2020.11.13.381871

**Authors:** Zhengfeng Lai, Runlin Guo, Wenda Xu, Zin Hu, Kelsey Mifflin, Charles DeCarli, Brittany N. Dugger, Sen-ching Cheung, Chen-Nee Chuah

## Abstract

Neurodegenerative disease pathologies have been reported in both grey matter (GM) and white matter (WM) with different density distributions, an automated separation of GM/WM would be extremely advantageous for aiding in neuropathologic deep phenotyping. Standard segmentation methods typically involve manual annotations, where a trained researcher traces the delineation of GM/WM in ultra-high-resolution Whole Slide Images (WSIs). This method can be time-consuming and subjective, preventing the analysis of large amounts of WSIs at scale. This paper proposes an automated segmentation pipeline combining a Convolutional Neural Network (CNN) module for segmenting GM/WM regions and a post-processing module to remove artifacts/residues of tissues as well as generate XML annotations that can be visualized via Aperio ImageScope. First, we investigate two baseline models for medical image segmentation: FCN, and U-Net. Then we propose a patch-based approach, ResNet-Patch, to classify the GM/WM/background regions. In addition, we integrate a Neural Conditional Random Field (NCRF) module, ResNet-NCRF, to model and incorporate the spatial correlations among neighboring patches. Although their mechanisms are greatly different, both U-Net and ResNet-Patch/ResNet-NCRF achieve Intersection over Union (IoU) of more than 90% in GM and more than 80% in WM, while ResNet-Patch achieves 1% superior to U-Net with lower variance among various WSIs. ResNet-NCRF further improves the IoU by 3% for WM compared to ResNet-Patch before post-processing. We also apply gradient-weighted class activation mapping (Grad-CAM) to interpret the segmentation masks and provide relevant explanations and insights.

## I. Introduction

**A**lzheimer’S disease is the sixth leading cause of death (122,019 deaths in 2018) in the United States and the number of people with Alzheimer’s disease is predicted to increase to 13.8 million in the United States by mid-century [1]. To analyze the progression of this disease, neuropathologists assess postmortem brain tissue slides, where they identify diverse and subtly-differentiated morphologies that are important for the diagnosis of Alzheimer’s disease [2]. Traditionally, neuropathologists compare these morphologies by studying brain tissue slides using microscopes, and the pathology workforce is dwindling, and items are needed to augment the ability of the pathologist [3]. Recently, with the help of digital slide scanners, details could be preserved by scanning physical tissue slides into ultra-high resolution whole slide images (WSIs) so that trained personnel are able to identify and analyze morphologies by viewing these WSIs [4], [5] through select software (such as Aperio ImageScope and QuPath [6]).

There are many pathologies within the brain that define neurodegenerative diseases, such as Alzheimer’s disease, and their locations can be extremely important to gain insights into disease pathophysiology. One hallmark pathological feature of Alzheimer’s disease is the presence of extracellular Amyloid-*β* plaques in human’s brain [7], [8]. Most plaques are found in grey matter (GM) while some have also been reported in white matter (WM) [9]. As such, an important task for assessing neurodegenerative disease pathologies (such as Alzheimer’s disease [2], [10]) is to dichotomized between GM and WM. Furthermore, incorporation of region of interest detection algorithms, such as GM/WM segmentation will be highly advantageous in quantitative pathological studies providing further insights into selective vulnerability and disease pathophysiology. However, manual segmentation of these ultra-high resolution WSIs can be time-consuming and not a cost-effective means for large scale projects. Moreover, manual segmentation could be subjective and have inter-rater reliability issues. Fig. 1 shows two examples, in each of which two trained personnel draw considerably different boundaries between GM and WM in the same brain WSIs.

**Fig. 1.**
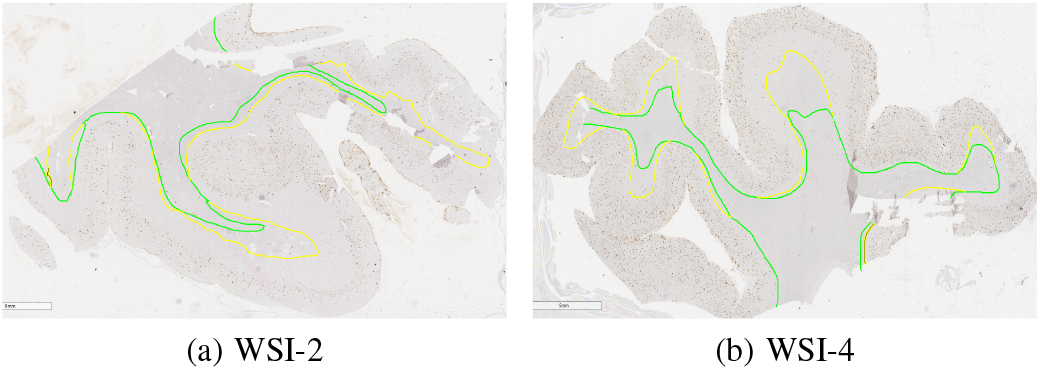
WSIs annotated by two trained personnel. (Green and yellow colors denote the two independent annotations.)

Therefore, there is a diverse need to build automated and efficient segmentation algorithms that are generalizable enough to apply to dataset of WSIs to provide a scalable means for deeper phenotyping of neurodegenerative diseases. There are many proposed methods in both traditional image processing and deep learning for WSI segmentation. However, these have been applied to the tissue slides primarily from other parts of human body (tongue, breast, lymph node, rectum [11], and skin [12]), anatomical brain (instead of tissue level) [13], or images reconstructed from magnetic resonance imaging (MRI) and computerized tomography (CT) [14], [15].

Although these methods are helpful in the analysis of WSIs in various applications, there are several issues that limit the direct applications of these methods to our WSIs for the analysis of Alzheimer’s disease. First, there are some unwanted artifacts in WSIs, such as tissue residues, tissue folds, bubbles, and/or dust. A sample of these artifacts is shown in Figure 2. These artifacts set obstacles for segmentation methods to separate the background out accurately. Second, our WSIs are scanned at ultra-high resolution to retain cellular level medical details (at 20x magnification) and thus the average resolution of these WSIs exceeds 50, 000 by 50, 000 pixels, making the files very large and difficult to process in a timely manner. To solve this resolution issue, current methods can be divided into two categories: either downsample original images of ultra-high resolution or divide original images into small patches for separate processing [16]. To provide a proof of concept for a workflow for GM/WM segmentation algorithm and to minimize the need for manual segmentation and tracings, we utilize 30 WSIs with annotations, far less than previous datasets used in [16]. Therefore, downsampling is not applicable for deep learning based methods. Both the plaques and cells with sizes ranging from 20 × 20 to 100 × 100 in WSIs are distinguishable features of GM and WM while simple downsampling may make them less clear or even disappear [17], which results in their limited performance. We need to preserve the distributions and characteristics of plaques in WSIs for the study of Alzheimer’s disease. Therefore, in this paper we aim to focus on the patch-based approach.

**Fig. 2.**
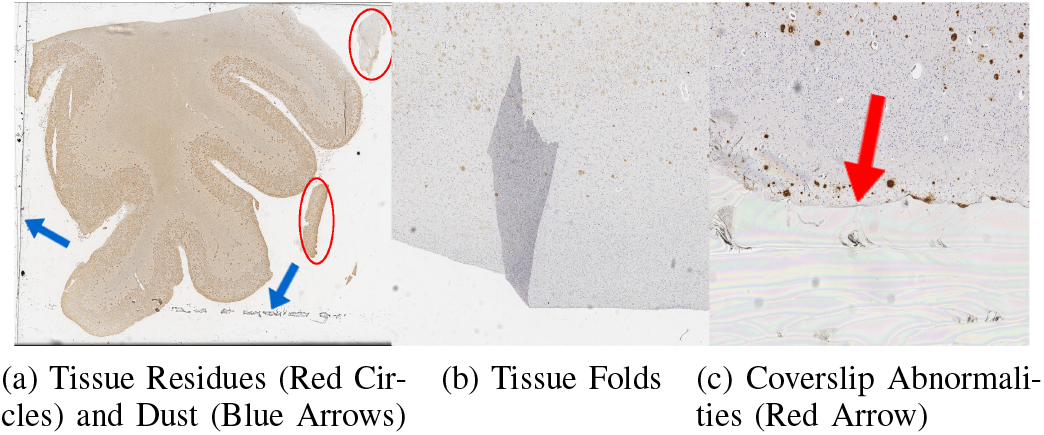
Sample Artifacts in WSIs.

In this paper, we propose an entire pipeline that incorporates CNN modules used for segmenting GM/WM regions with postprocessing module that removes artifacts and residues, as well as generates XML annotations that can be displayed on original WSIs. We implement two baseline CNN models that are widely deployed in medical image segmentation problems, namely a fully convolutional network (FCN) [18] and a U-Net model [19], to perform pixel-wise segmentation of GM and WM. Then, we propose ResNet-Patch, a novel mechanism that transforms the pixel-wise segmentation problem of ultra-high-resolution WSIs into a patch-wise classification problem in the first stage, then subsequently converts it back to the original segmentation task according to various resolution requirements of outputs. ResNet-Patch is built-upon our prior work applying CNN to GM and WM segmentation problem [17] with a limited dataset (18 WSIs from Alzheimer’s disease cases). Although our initial results were promising, the output segmentation mask was not desirable in terms of accuracy due to the presence of residues and artifacts. ResNet-Patch addresses these challenges by incorporating post-processing pipelines to reduce artifacts and training the network with heterogeneous input (WSIs from cases with and without Alzheimer’s disease). Besides, to solve the prediction inconsistency issue in the first stage of ResNet-Patch, we incorporate ResNet-Patch with a Neural Conditional Random Field (NCRF) module to model the spatial relationships among neighboring patches, referred to as ResNet-NCRF.

Our contributions can be summarized as follows:

- We propose an automated Amyloid-*β* WSI segmentation pipeline that includes both Convolutional Neural Network (CNN) for separating GM and WM, and post-processing used for removing artifacts as well as generate XML annotations that can be visualized and displayed on original-size WSIs via Aperio ImageScope
- For the segmentation of GM and WM in gigapixel WSIs, we propose ResNet-Patch that is memory-efficient and can achieve superior GM/WM Intersection over Union (IoU) compared to U-Net and FCN. Furthermore, we integrate the NCRF module that incorporates spatial correlations among neighboring patches into ResNet-Patch. This is a novel application of CNN based segmentation techniques to this medical problem.
- We apply Gradient-weighted Class Activation Mapping (Grad-CAM) [20] to illustrate what CNN learns and relies on to distinguish GM from WM and confirm they are explainable from a pathologic perspective by consulting a domain expert in neuropathology.

Our results show the U-Net pipeline can segment GM and WM with IoU of 91.52% and 82.02% respectively when compared to expert annotations while the ResNet-Patch pipeline achieves 93.06% IoU on GM and 83.80% IoU on WM after post-processing. ResNet-NCRF can achieve 84.27% IoU on WM without post-processing. All of them are much higher than FCN that can only achieve 77.39%/57.29% in GM/WM respectively. The rest of the paper is organized as follows: Section II discusses related work, Section III describes our dataset, Section IV introduces pipeline architectures, Section V presents our results, and Section VI concludes the paper.

## II. Related work

### A. Image Processing-Based Segmentation

Tissue segmentation can be a critical step in the disease diagnosis [21]. Many traditional image processing algorithms have been designed for whole slide images (WSIs) segmentation. Hiary et al. [22] built an automatic algorithm based on k-means clustering that works on pixel intensity and texture features. Bug et al. [12] applied global thresholding at the mean value of the Gaussian blurred Laplacian of the greyscale image. In [23], stain concentration and porospity analysis was applied to the segmentation of H&E stained WSIs. Although these methods are computational efficient, the segmentation results are very limited due to inter and intra-variations in staining and color contrast, resulting in general failure for the above mentioned methods on hold-out test sets [24]. Additionally these methods have only been tested on non-brain WSIs like breast, tongue and skin, so their applicability and feasibility on our brain WSIs has yet to be determined.

### B. Deep Learning-Based Segmentation

In recent years, deep learning methods have gained vast popularity in image segmentation problems. For example, Hiary et al. [14] proposed an automatic method that can segment brain tumor by using convolutional neural network (CNN) to extract local features and global contexts. In [25], Milletari et al. proposed Hough-CNN to perform segmentation of deep brain regions in MRI and ultra-sound images. Besides these CNN methods, FCN [11] and U-Net [24] based architectures are predominant choice in medical image segmentation problems [19], [26]. In this section, we will discuss the details of these two architectures that are used as benchmarks for this paper.

#### 1) FCN

Fully convolutional network (FCN), due to its arbitrary input size and better localization performance [27], was a popular choice in the recent works of medical image segmentation [28]. When applied to tissue segmentation of histopathological WSIs, FCN outperforms traditional methods and achieve less outliers and more stable results [11]. FCN is used for multi-organ segmentation [29], in both 2D slices of 3D volume [30] and 3D images [31]. Besides regular FCNs, modified Cascaded FCNs have been deployed in liver segmentation, where the first FCN performed Region of Interest (ROI) and second performed the lesion segmentation [32]. Multi-stream FCN can improve accuracy by allowing system to take multiple forms of the same organ [33]. Moreover, focal loss is applied on the FCN due to the class imbalances of training data. The number of false positives can be removed by applying the focal loss on the FCN network [34] [35].

The most recent FCN model (as shown in Fig. 3) we explored is based on the architecture and hyper-parameters in [11]. We used FCN-8s model from the original FCN paper [27], which combined predictions from the final layer, pool4 layer and pool3 layer at the stride of 8. The FCN architecture contains 7 convolutional layers. The first two layers have the filter size 5 × 5. The third and fourth layers have filter size 3 × 3. The fifth layer has the size of 11 × 11. The last two layers have 1 × 1 convolution, which are equivalent to the fully connected layers. The number of filters at each layer are 16, 32, 64, 64, 1024, 512 and 2 respectively as shown in Fig. 3. Filter size of 2 × 2 and stride of 2 are used to all the max pooling layers. The number of max pooling layers is based on original FCN paper [27], which are inserted after first five convolutional layers to reduce memory requirements of the network. The batch normalization layer and sigmoid activation layer are added after every convolutional layer for the regularization and convergence purposes. Drop out layer with drop out rate of 0.2 is added after every convolutional layer to reduce the probability of overfitting. The batch size is 128.

**Fig. 3.**
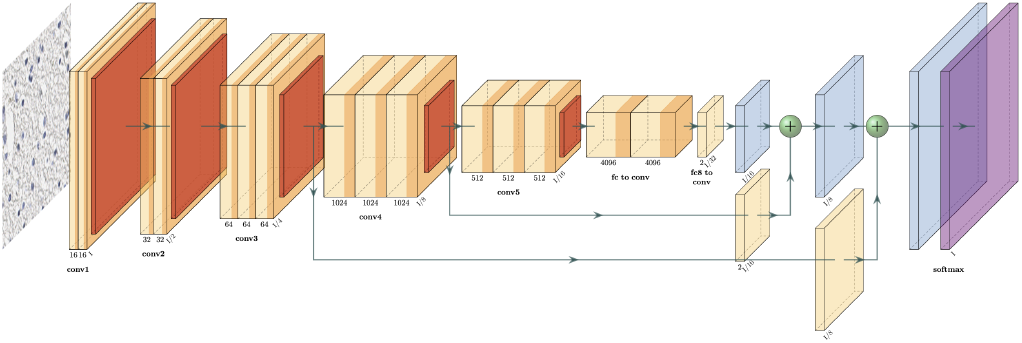
FCN Architecture.

By benchmarking the results of different loss functions, we chose the combination of the categorical cross entropy loss and focal loss for FCN [34] [35]. By setting the weight of different output classes to be 1 and training FCN on categorical cross entropy loss and then on focal loss, the average validation results of three output classes outperformed using either categorical cross entropy or focal loss alone (weighted or unweighted).

#### 2) U-Net

U-Net is one of the state-of-the-art models in medical image segmentation and has gained great popularity in different medical problems. For example, Dong et al. [36] built a fully automatic method based on U-Net for brain tumor segmentation of MRI. In [37], Rampun et al. proposed a 2D fetal brain segmentation of MRI by modifying the U-Net architecture. A major drawback of U-Net based methods is that they require computationally intensive operations on GPUs and take a great deal of processing power with very trivial gain in performance [38].

The updated U-Net model we investigated is based on the architecture and hyper-parameters in [24]. As shown in Fig. 4, the contracting path of the network has four convolution levels. Each level consists of two consecutive 5 × 5 convolution operations (zero-padded convolutions) with Exponential Linear Unit (ELU) [39] activation functions, followed by a 2 × 2 max-pooling operation with stride 2 for down-sampling. At each down-sampling level, the number of feature channels is doubled (32 − 64 − 128 − 256 − 512). The expansive part of the network also has four convolution levels. Each level consists of a 2 × 2 transposed convolution that halves the number of feature channels, a concatenation with the corresponding contracting level output, and two 5 × 5 convolutions with ELU activation. To speed up learning and provide some regularization effect, we used batch normalization [40] after each convolutional layer. We also incorporated drop-out [41] after the first convolutional layer at each level, both in the contracting and the expansive path. A final 1 × 1 convolution with 3 output channels is then used to map the last feature map to the class prediction output.

**Fig. 4.**
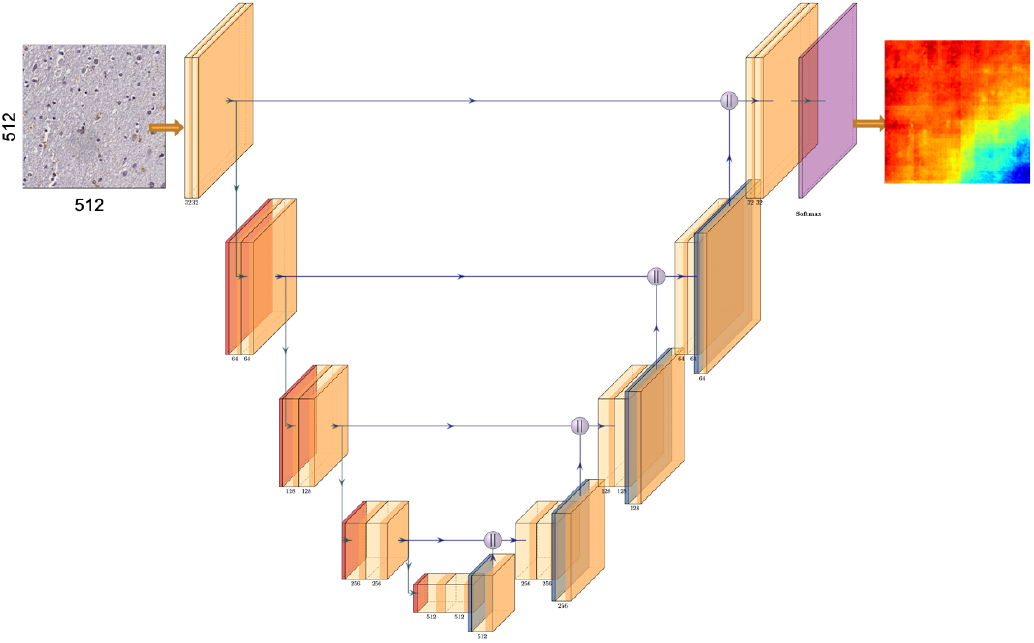
U-Net Architecture.

Prior to the training, all convolution kernels were initialized using *He’s uniform* initialization [42]. We used the *Adam* optimizer [43] with an initial learning rate of 0.001. The *α* parameter for the ELU activations was set to 1.0, the drop-out rate was 0.2, and the batch size was 16. To diminish the effect of class-imbalance issue in our dataset, we weighted the categorical cross entropy loss function by the inverse of class frequencies in the training dataset. However, there is another obvious limitation among these deep learning-based methods: these networks were specifically designed and tested on specific medical applications such as tumor detection or image modality (MRI that has standard image formats and CT), making it difficult to generalize existing findings to our human brain tissue WSIs that have no standard formats [44]. While MRI and CT are representations of a global view of the whole brain, our WSIs can only capture a local view of one region of brain, unlike semantic objects in natural images, there are subtle and gradual changes of pathological features rather than the sharp color edges in clinically meaningful regions Many CNN models rely heavily on annotations, indicating the difficulty for them to self-identify residues and artifacts in the tissue slides if the annotations are not provided. For example, in [17], although their CNN architecture is able to distinguish GM from WM, it is unable to remove residues and artifacts since the corresponding annotations are not provided.

### C. Patch-based Approach for Ultra-high Resolution Images

Most of these deep learning methods have only been tested on images with low to medium resolution of up to a few megapixels. Chen et al. [16] regarded images with resolution up to 30 million pixels as ultra-high resolution images. They proposed a method that integrates both global downsampled images and local patches while they only tested their methods in images with resolution up to 30 million pixels. However, the average resolution of our WSIs exceeds 2 gigapixels. Downsampling used in [16] cannot be directly transferred to our case as we have only 30 WSIs with proper annotations, which indicates the shortage of training data, while the datasets used in their experiments consist of more than 2,000 images. Besides, we need to preserve minute details of pathologies (such as plaques) and cells in our WSIs while downsampling will unavoidably lose these important features. In [17], the authors conducted their experiments by introducing downsampling in the training process, which resulted in their limited performance on final masks.

Because of the ultra-high resolution and details that needs to be preserved, recently many studies applied patch-based methods to WSIs [45]–[47]. They extracted small patches (e.g. 256 × 256 pixels) from WSIs and then trained a deep CNN model. Although these studies solved the resolution issue, most of these applications are classification problems (e.g. breast cancer classification [47]) while our problem is a segmentation problem. Besides, as the patches were extracted and trained independently in these studies, the spatial correlations shared by small patches and their neighbors may not be modeled explicitly. As a result, the predictions over neighboring patches may not be consistent and there may exist isolated outliers in the patch-level probability map during inference time [48], [49]. To incorporate the spatial correlations between neighboring patches, authors in [48] used Conditional Random Field (CRF) to model spatial correlations and refine the predicted probability map in the postprocessing stage. However, due to the high computational overhead, the authors can only feed five features from patch representations into the CRF inference algorithm. In this paper, we propose a pipeline that first converts the GM/WM segmentation to a patched-based classification problem, and then generates pixel-level segmentation output based on the classification results. Subsequently, we propose a one-stage segmentation method that incorporate spatial correlations among neighboring patches through a Neural Conditional Random Field (NCRF) [50] layer.

## III. Datasets

### A. Overview

Our dataset consists of 30 Whole Slide Images (WSIs) stained with an Amyloid-*β* antibody (4GB, recognizing residues 17–24, dilution 1:1600, BioLegend (formally Covance) catalog number SIG-39200) [51] from the temporal cortex. As these WSIs were digitized by Aperio AT2 at up to 40 × magnification, the resolution is nearly 60, 000 50, 000 pixels each on average. These 30 WSIs can be split into two sets: one set includes 18 cases (10 males and 8 females) that had a pathologic diagnosis of Alzheimer’s disease with an average age at death of 84 ± 7 years while the other set (12 cases) lacked a pathologic diagnosis of Alzheimer’s disease (Non-Alzheimer’s disease, NAD). Of NAD cases, 5 had a diagnosis of cerebrovascular disease, and 1 with metastatic carcinoma. The Ethnoracial make up of the cohort was 3 Hispanics (10%), 5 African Americans (17%), and 22 non-Hispanic White (73%) descendants.

These studies utilized tissues only from human post-mortem. Only living subjects are confirmed as Human Subjects under federal law (45 CFR 46, Protection of Human Subjects). All participants or legal representative approved informed consent during the life of the participant as part of the University of California Davis Alzheimer’s disease Center program. The data collection process followed current laws, regulations and IRB guidelines. All of these 30 WSIs have been de-identified, which do not have personal health information like names, social security numbers, addresses, and phone numbers and are shared with a randomly generated pseudo-identification number. To further protect data confidentiality, we name the AD cases as WSI-1 to WSI-18 and NAD cases as WSI-19 to WSI-30.

### B. WSI Analysis

To further characterize the datasets, we compare the size of GM/WM region, the ratio between the tissue and the whole slide, and the ratio between GM/WM region and the whole tissue, separately in both AD set (18 WSIs) and NAD set (12 WSIs). Fig. 5 displays the box plot charts for visualizing these comparisons where differences between AD set and NAD set are noticeable: the average absolute size of WM of AD cases is around 57.35% larger than that of NAD cases (Fig. 5b), so as the percentage of WM in tissue as shown in Fig. 5a; the thickness ratio between GM and WM in NAD cases tends to be nearly 53.53% larger than that in AD cases (Fig. 5c). Also, the average for NAD sulci is around 2 per slide while this is nearly 4 per slide for AD cases.

**Fig. 5.**
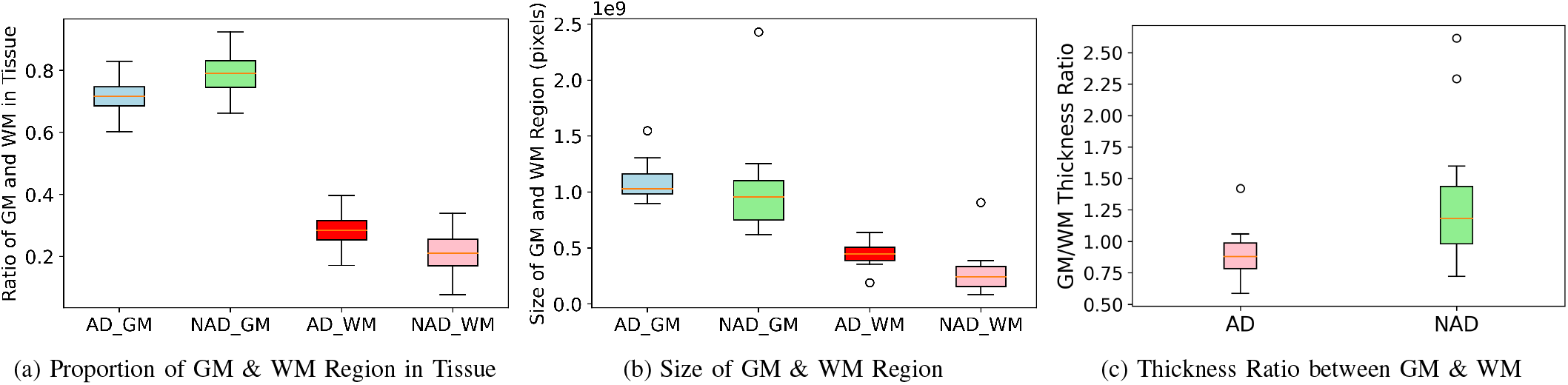
Dataset feature comparison. Box plot chart comparing (a) the proportion of GM/WM region in tissue for both AD cases and NAD cases, respectively, (b) the absolute size of GM and WM, and (c) the thickness ratio between GM and WM in AD and NAD cases.

### C. Training Data Preparation

To deal with the issue of ultra-high resolution, we use Pyvips Library [52] to set up image processing pipelines on original-resolution WSIs instead of directly manipulating these slides. Hence, we avoid loading the entire image into memory at once for processing because Pyvips can stream the image in parallel from the first step to the last step of pipelines simultaneously. The proposed approach in [17] manually selected regions from GM, WM and background and cropped small tiles from these regions separately. This manual action could result in limited variety of datasets and involve human interventions. As we have pixel-wise annotations on GM and WM for all 30 WSIs, we randomly cropped patches 256 × 256 and labeled them as follow. Each patch is labeled with the category of the central pixel of that patch tracing back to our pixel-wise ground truth.

The datasets for FCN and U-Net are the same-split of the 30 WSIs but cropped into 512 × 512 non-overlapping patches.

In our experiments, 20 WSIs (12 AD cases and 8 NAD cases) were randomly selected for training and validation while the remaining 10 WSIs (6 AD cases and 4 NAD cases) were used for hold-out testing and inference. These WSIs were annotated by trained personnel (K.M and B.D) at pixel level.

## IV. Methodology

This section introduces our proposed pipeline for the segmentation of GM and WM. As shown in Fig. 6, we generate the masks first via the CNN module, subsequently we use the post-processing module to remove artifacts, finally we display the mask on the original-size WSI to generate XML annotations as the final output, where yellow contours denote the boundary between GM and WM while cyan contours segment the tissue from the background. In the CNN module, we investigate FCN and U-Net first, then propose ResNet-Patch and ResNet-NCRF and compare their performance.

**Fig. 6.**
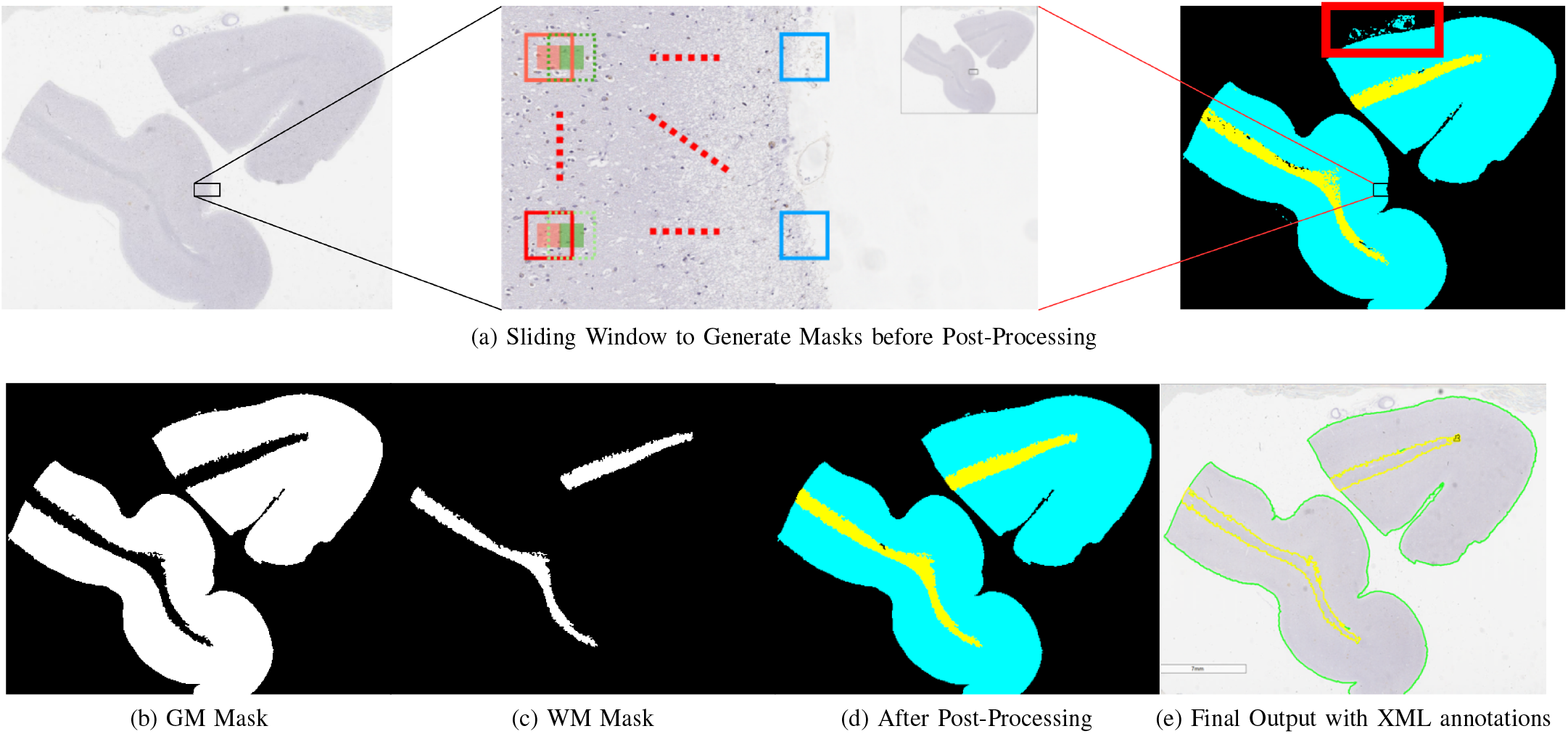
Segmentation Pipeline and Output of XML annotations.

### A. ResNet-Patch

ResNet-Patch consists of two stages: transform the problem to patch-based classification and convert back to pixel-wise segmentation of WSIs.

#### 1) Patch-based Classification

In this stage, we transform the pixel-level segmentation problem into a patch-based classification problem. For image classification tasks, a myriad of different CNN architectures have been proposed, such as AlexNet, VGG, ResNet [53] in the past few years. While design motivations of each architecture are greatly different, there is increasing evidence showing that the features extracted from these state-of-the-art architectures are quite akin and their improvements gradually become trivial and start to converge [54]. To produce reasonable results with relatively minimized complexity, we decide to select a simple architecture to pursue the tradeoff between complexity and performance. In [55], He et al. showed their ResNet architecture has fewer filters and lower complexity but achieves similar performance compared to other state-of-the-art architectures. As such, we select ResNet-18 as our backbone network and the fundamental basis of our methods.

We modified ResNet-18 by redefining the last fully connected layer to output three categories: GM, WM, and the background of tissue slides. As we transform the segmentation problem into a classification problem in this stage, the function of ResNet-18 here is to extract the features from each patch and classify its corresponding category. We adapted pre-trained parameters except for the last layer from ResNet-18 trained on ImageNet because it has already shown the ability of extracting useful features from natural images. We trained this modified ResNet-18 for 10 epochs. We used *Adam* optimizer [43] and the initial learning rate is 0.001 for the first five epochs and 0.0001 for the remaining 5 epochs. The batch size was set as 16. For the loss function, considering the effect of class-imbalance issue, we introduced categorical weights to the cross entropy loss based on the inverse of class frequencies in our training set.

#### 2) Pixel-wise Segmentation

In this stage, we utilize the results from the classification task to construct pixel-wise segmentation output. In the previous stage, each patch is classified to a corresponding output class (GM, WM, or background) using our modified ResNet-18 model.

In order to achieve pixel-wise segmentation results, we use a sliding window approach to extract each patch until the whole image is fully covered. And the resolution of output masks will be decided by the step size. For example, as shown Fig. 6a, step size here is set as 128 while the patch size is 256. The red patch in Fig. 6a is fed into our modified ResNet-18 first, subsequently receives the prediction category. After that, if output masks are required to be the same size of original WSIs, all pixels in the central area (red block) with the size of step size × step size (128 × 128 here) will be classified as the same category predicted by the modified ResNet-18 model. Then we make a step forward with the step size to the green patch with dotted line and repeat the same action, so the green block next to the red block will be labeled with the same category. By repeating this action, after the sliding window traverses the whole image, we can get segmentation masks for GM, WM (as shown in Fig. 6b, 6c) and background separately. As such, we can see the accuracy of output masks is determined by step size: if step size is larger, outputs will be less accurate around boundaries but inference complexity will be reduced; if step size is smaller, outputs will be more accurate but inference complexity is also increasing. Hence, ResNet-Patch provides flexibility to achieve different trade-offs between performance and complexity.

### B. ResNet-NCRF

This section introduces the extension of ResNet-Patch, named as ResNet-NCRF, which uses the same mechanism and backbone as ResNet-Patch while integrating a Neural Conditional Random Field (NCRF) module that incorporates spatial relationships among neighboring patches. ResNet-NCRF (as shown in Fig. 7) consists of two components: ResNet-18 component is used to extract features and encode each patch as an embedding (a vector representation with a fixed length) by taking a grid of patches as input; NCRF component is used to model spatial correlations among the grid of patches and output the probability of central patch in the grid.

**Fig. 7.**
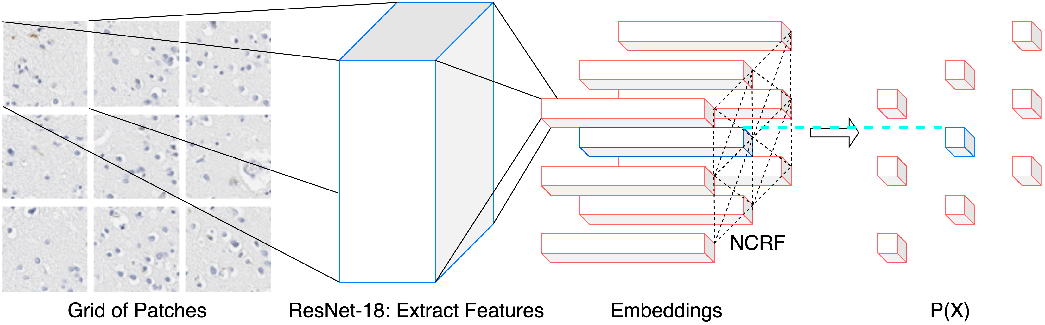
NCRF Architecture.

#### 1) Spatial Correlations with NCR

Given a grid of patches, we define a random field *X* = {*X*_1_, *X*_2_, …, *X*_*N*_} as the random variables related to each patch, where *N* is the number of patches in the grid, e.g. 9 for a grid of 3 × 3. Each random variable represents a label from the set of {*GM, WM, background*}, conditioned on observations *I* ={*I*_1_, *I*_2_, …, *I*_*N*_}, where *I* is the embedding of each original patch extracted by the CNN component. This set of label is defined as *x* ={*x*_1_, *x*_2_, …, *x*_*N*_}. Therefore, different from the CRF methods for pixel labeling, *I* is not a set of pixel attributes such as RGB color values or intensity but a set of patch descriptors: CNN features from each patch. The distribution of (*X, I*) can be a CRF if the random variables of set *X* conditioned on observations *I* meet the requirements of Markov property. Hence the structured prediction can be transformed as:

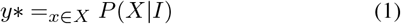

Where P(X | *I*) is defined as a Gipps distribution:

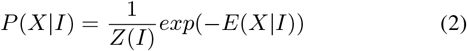

In a fully-connected CRF [56], the energy of Gibbs can be written as:

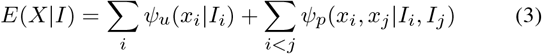

where *i, j* ∈ {1, 2, …, *N*}. The energy function *E*(*X*|*I*) is the summation of two terms: a unary potential *ψ*_*u*_(*x* |*I*) and the pairwise *ψ*_*p*_(*x*_*i*_, *x*_*j*_ *I*_*i*_, *I*_*j*_). The unary potential can be considered an initial estimate of the cost of patch *i* with its label *x*_*i*_ given its corresponding embedding *I*_*i*_. Similarly, the pairwise potential estimates the joint cost of patch *i, j* with their label *x*_*i*_, *x*_*j*_ given their corresponding embeddings *I*_*i*_, *I*_*j*_. In other words, the pairwise potential is a measure of relationships between two patches and is defined over every pair of patches in the grid. In [57], they use the cosine similarity to define the pairwise potential as:

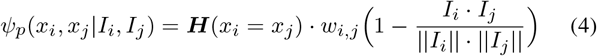

where ***H***(*I*_*i*_ = *I*_*j*_) is an indicator function that checks the label compatibility between *X*_*i*_, *X*_*j*_. By using the cosine similarity, the spatial correlations between a pair of patches can be modelled by encouraging lower cost in the case that *X*_*i*_, *X*_*j*_ are assigned with the same label if the embeddings *I*_*i*_, *I*_*j*_ are similar. *w*_*i,j*_ is a single, trainable parameter which the authors claimed related to spatial distance between patches [57].

#### 2) End-to-End Back-propagation

To compute the cross-entropy loss with the ground truth labels, we need to get the marginal distribution of each patch label *x*_*i*_ so that the standard back-propagation algorithms can be applied here to achieve end-to-end training. A mean-field approximation method was proposed in [58] to approximate maximum posterior marginal inference. As exact marginal inference is intractable, by using the mean-field approximation, the original CRF distribution *P* (*X*) can be transformed to *Q*(*X*) which is obtained by the product of independent marginal distributions, as shown in (5):

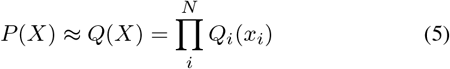

Here *Q*(*X*) is a simpler distribution compared to *P* (*X*). The 𝕂L divergence between *Q*(*X*) and *P* (*X*) is shown in (6).

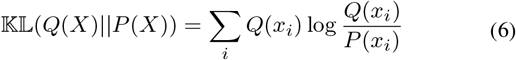

The first item is the negative of entropy while the second item is cross-entropy. Assuming 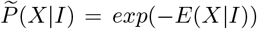 as the unnormalized CRF distribution, we can derive the cross-entropy item as shown in (7).

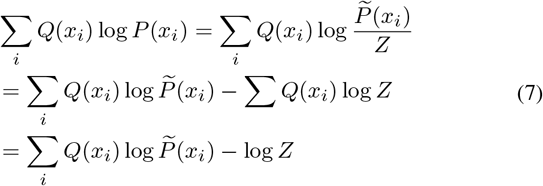

For minimizing 𝕂𝕃 divergence, −log *Z* can be dropped as this is an additive constant. This leaves us with the following optimization problem:

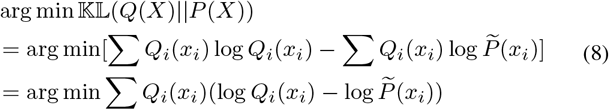

As 𝕂𝕃 divergence cannot be negative, the minimum of Equation 8 can be derived as:

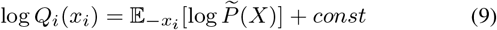

This is the update equation for each marginal distribution *Q*_*i*_(*x*_*i*_), where 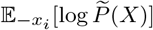 refers to the expectation of 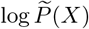 on all *x* except *x*_*i*_.

In summary, our mean-field marginal inference algorithm is shown in Algorithm 1.

### C. Post-processing

We design a post-processing approach to refine our segmentation output masks and remove tissue residues in our WSIs. The full-resolution predicted outputs are down-sampled by a factor of 16 (1/4 in width and 1/4 in height) to reduce computational complexity. Two consecutive area openings are applied to remove fuzzy predictions of GM and WM with pixel area *<* 20, 000 and an area closing is applied to remove fuzzy prediction of background with pixel area *<* 12, 500. Then, small tissue residues with area smaller than 5% of the WSI area are removed. In the end, a morphological opening with a disk-shaped kernel with radius of eight is applied to smooth the boundary and the output was up-sampled back to the original resolution. The kernel sizes used for morphological opening and the size threshold for pixel areas to be removed are obtained empirically. Fig. 6a, 6d is an example showing the difference between before and after we apply this post-processing to remove tissue residues.

#### Algorithm 1

Mean-filed Marginal Inference Algorithm

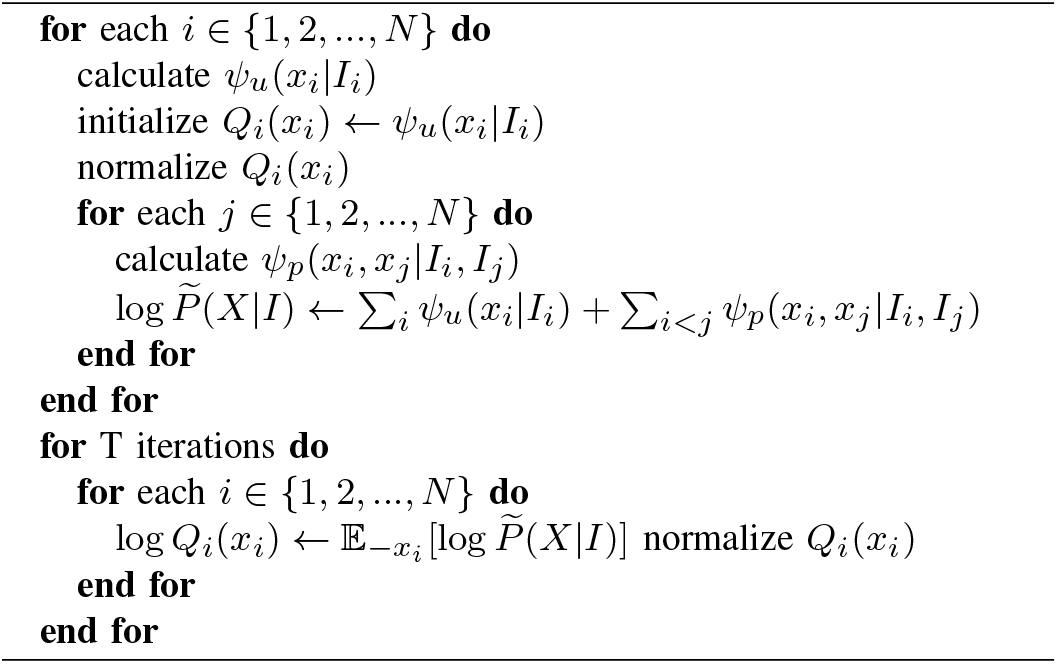

In addition to these steps, we also generate XML annotation files of GM and WM segmentation boundaries by finding contours of GM and WM segmentation masks respectively as shown in Fig. 6e. These XML files are in the same format of our ground truth annotations and can be visualized and displayed on original-size WSIs using Aperio ImageScope. They are also downsampled by 50× to smooth out the boundary for faster visualization.

## V. Results

### A. Quantitative Results

#### 1) Pixel-wise Classification

To measure the pixel-wise classification performance of these three methods, we select different indexes as follows: Accuracy (10), Recall (11), Precision (12) and F1-Score (13).

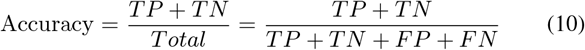

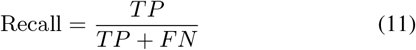

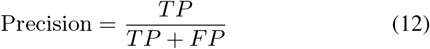

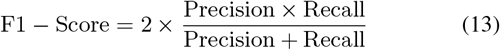

The results on hold-out testing set are summarized in Fig. 8. We can see the average F1-Score of GM is higher than that of WM in all these four methods, the same as Recall and Precision, while the average Accuracy of GM is slightly lower than that of WM. ResNet-Patch has 1% higher F1-Score than U-Net in both GM and WM while ResNet-NCRF further improves the performance. FCN is nearly 23% worse than ResNet-Patch/ResNet-NCRF in F1-Score mainly due to its limited performance on Precision. Both ResNet-Patch/ResNet-NCRF and U-Net achieves 95% in terms of F1-Score in GM and slightly lower, around 90%, in WM.

**Fig. 8.**
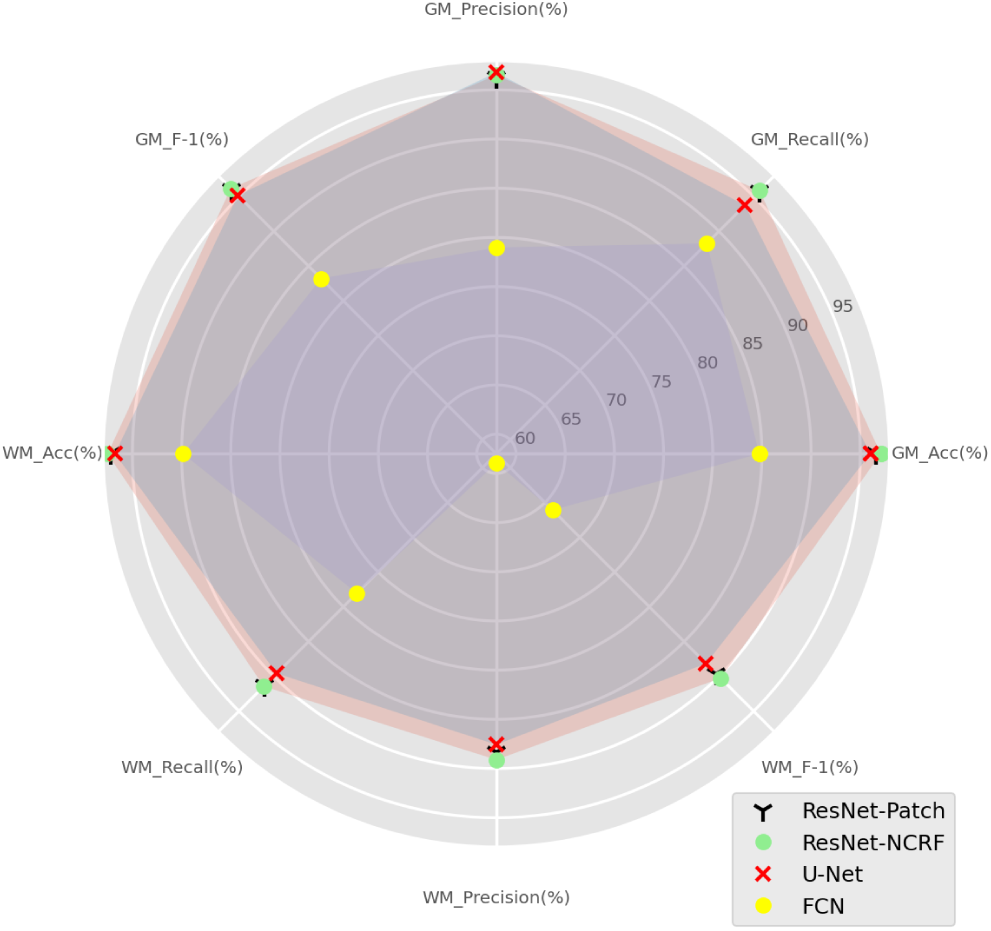
Pixel-wise classification accuracy comparison.

**TABLE I.**
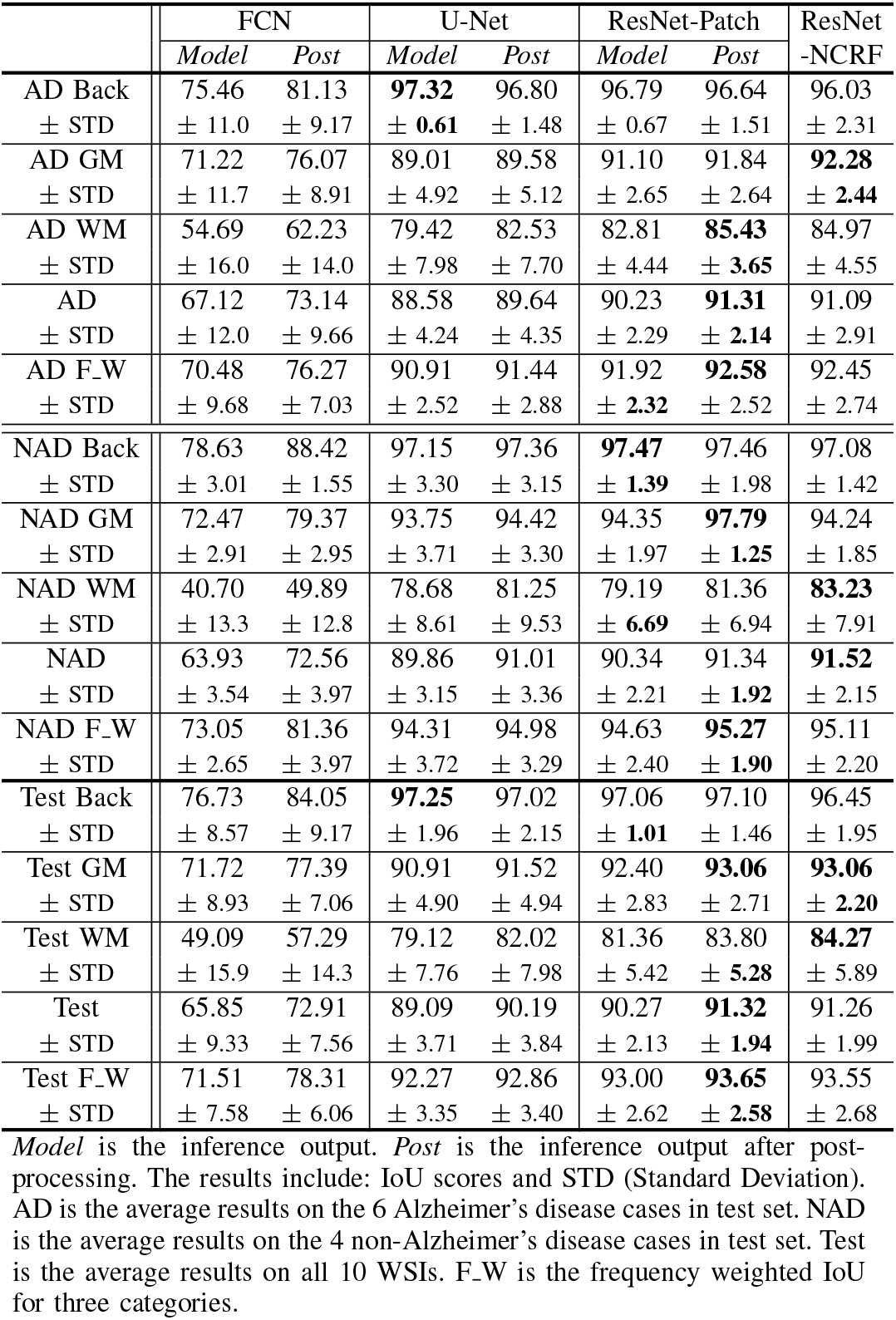
Pixel-wise IoU/STD Comparison FOR AD, NAD, AND OVERALL HOLD-OUT TEST SET

|  | FCN |  | U-Net |  | ResNet-Patch |  | ResNet-NCRF |
| --- | --- | --- | --- | --- | --- | --- | --- |
|  | *Model* | *Post* | *Model* | *Post* | *Model* | *Post* |  |
| AD Back | 75.46 | 81.13 | **97.32** | 96.80 | 96.79 | 96.64 | 96.03 |
| ± STD | ± 11.0 | ± 9.17 | ± **0.61** | ± 1.48 | ± 0.67 | ± 1.51 | ± 2.31 |
| AD GM | 71.22 | 76.07 | 89.01 | 89.58 | 91.10 | 91.84 | **92.28** |
| ± STD | ± 11.7 | ± 8.91 | ± 4.92 | ± 5.12 | ± 2.65 | ± 2.64 | ± **2.44** |
| AD WM | 54.69 | 62.23 | 79.42 | 82.53 | 82.81 | **85.43** | 84.97 |
| ± STD | ± 16.0 | ± 14.0 | ± 7.98 | ± 7.70 | ± 4.44 | ± **3.65** | ± 4.55 |
| AD | 67.12 | 73.14 | 88.58 | 89.64 | 90.23 | **91.31** | 91.09 |
| ± STD | ± 12.0 | ± 9.66 | ± 4.24 | ± 4.35 | ± 2.29 | ± **2.14** | ± 2.91 |
| AD F.W | 70.48 | 76.27 | 90.91 | 91.44 | 91.92 | **92.58** | 92.45 |
| ± STD | ± 9.68 | ± 7.03 | ± 2.52 | ± 2.88 | ± **2.32** | ± 2.52 | ± 2.74 |
| NAD Back | 78.63 | 88.42 | 97.15 | 97.36 | **97.47** | 97.46 | 97.08 |
| ± STD | ± 3.01 | ± 1.55 | ± 3.30 | ± 3.15 | ± **1.39** | ± 1.98 | ± 1.42 |
| NAD GM | 72.47 | 79.37 | 93.75 | 94.42 | 94.35 | **97.79** | 94.24 |
| ± STD | ± 2.91 | ± 2.95 | ± 3.71 | ± 3.30 | ± 1.97 | ± **1.25** | ± 1.85 |
| NAD WM | 40.70 | 49.89 | 78.68 | 81.25 | 79.19 | 81.36 | **83.23** |
| ± STD | ± 13.3 | ± 12.8 | ± 8.61 | ± 9.53 | ± **6.69** | ± 6.94 | ± 7.91 |
| NAD | 63.93 | 72.56 | 89.86 | 91.01 | 90.34 | 91.34 | **91.52** |
| ± STD | ± 3.54 | ± 3.97 | ± 3.15 | ± 3.36 | ± 2.21 | ± **1.92** | ± 2.15 |
| NAD F.W | 73.05 | 81.36 | 94.31 | 94.98 | 94.63 | **95.27** | 95.11 |
| ± STD | ± 2.65 | ± 3.97 | ± 3.72 | ± 3.29 | ± 2.40 | ± **1.90** | ± 2.20 |
| Test Back | 76.73 | 84.05 | **97.25** | 97.02 | 97.06 | 97.10 | 96.45 |
| ± STD | ± 8.57 | ± 9.17 | ± 1.96 | ± 2.15 | ± **1.01** | ± 1.46 | ± 1.95 |
| Test GM | 71.72 | 77.39 | 90.91 | 91.52 | 92.40 | **93.06** | **93.06** |
| ± STD | ± 8.93 | ± 7.06 | ± 4.90 | ± 4.94 | ± 2.83 | ± 2.71 | ± **2.20** |
| Test WM | 49.09 | 57.29 | 79.12 | 82.02 | 81.36 | 83.80 | **84.27** |
| ± STD | ± 15.9 | ± 14.3 | ± 7.76 | ± 7.98 | ± 5.42 | ± **5.28** | ± 5.89 |
| Test | 65.85 | 72.91 | 89.09 | 90.19 | 90.27 | **91.32** | 91.26 |
| ± STD | ± 9.33 | ± 7.56 | ± 3.71 | ± 3.84 | ± 2.13 | ± **1.94** | ± 1.99 |
| Test F.W | 71.51 | 78.31 | 92.27 | 92.86 | 93.00 | **93.65** | 93.55 |
| ± STD | ± 7.58 | ± 6.06 | ± 3.35 | ± 3.40 | ± 2.62 | ± **2.58** | ± 2.68 |

*Model* is the inference output. *Post* is the inference output after post-processing. The results include: IoU scores and STD (Standard Deviation). AD is the average results on the 6 Alzheimer's disease cases in test set. NAD is the average results on the 4 non-Alzheimer's disease cases in test set. Test is the average results on all 10 WSIs. F.W is the frequency weighted IoU for three categories.

### 2) Pixel-wise IoU and Standard Deviation

Since our goal is pixel-level GM/WM segmentation, we use a well-known metric — Intersection over Union (IoU) to compare segmentation masks from different methods. We generate masks of GM, WM (as shown in Fig. 6b, 6c) and background for each WSI separately for different methods. Table. I summarizes IoU for the four CNN-based models we investigated. We calculate the average IoU of three categories in AD cases and NAD cases separately, as well as the mean value of three categories’ IoU and the frequency weighted IoU.

The slides shown in Fig. 2, Fig. 6, and Fig. 10 clearly indicate the inherent heterogeneity of brain tissues among different WSIs, where GM/WM regions have a great variety of sizes and shapes. Considering its variety of shapes, we also select standard deviation to describe how consistent and robust our methods are on different hold-out WSIs. Larger standard deviations indicates more performance inconsistency among heterogeneous slides. FCN achieves inferior performance compared to U-Net (its mean IoU is more than 20% lower than ResNet and U-Net), which is similar to the results reported in [59], [60]. Hence we will only compare ResNet-Patch/ResNet-NCRF with U-Net in the subsequent sections.

From Table. I, ResNet-Patch/ResNet-NCRF achieves the highest IoU and lowest standard deviation in the majority of indexes, which indicates that ResNet-Patch/ResNet-NCRF is more stable and robust to a new hold-out test set compared to FCN and U-Net. Specifically, the mean IoU of ResNet-Patch achieves more consistent segmentation results (less uncertainty) for both AD and NAD cases, since the difference between that of AD and that of NAD is only around 0.1% while there is more disparity in the confidence interval for AD vs. NAD cases for U-Net. The increased uncertainty in the segmentation results for AD compared to NAD is expected due to the existence of large amounts of plaques in AD cases.

Table. I shows that ResNet-Patch achieves 1.6% higher than U-Net in the mean IoU of AD cases, with smaller confidence intervals. Both ResNet-Patch and U-Net achieve compatible IoU for NAD cases, but there is less uncertainty (smaller confidence interval) for ResNet-Patch. Therefore, we can conclude that ResNet-Patch shows stronger distinguishing ability in GM and WM among AD cases where boundaries are harder to distinguish due to pathologies associated with AD. ResNet-NCRF further improves ResNet-Patch’s ability in WM among NAD cases and removes the need for post-processing.

Our post-processing can achieve more than 1% of improvement in IoU for both ResNet-Patch and U-Net and is able to visually remove most of the tissue residues successfully as shown in Fig. 2e. Table. I shows that ResNet-NCRF (without any post-processing step) can achieve comparable or superior performance compared to ResNet-Patch with post-processing. As ResNet-NCRF alone could generate smooth masks with few noisy pixels, post-processing module is not needed for ResNet-NCRF and the improvement is trivial. We decide not to incorporate the post-processing module into ResNet-NCRF. Hence, we may conclude the NCRF module is efficient in capturing spatial correlations among patches to increase the prediction consistency during inference.

#### 3) Training Process Comparison

To compare the robustness of ResNet-Patch/ResNet-NCRF with U-Net, we also analyze the characteristics of training process of ResNet-Patch/ResNet-NCRF and U-Net as shown in Fig. 9. During our experiments, we find the epoch loss of U-Net on the validation set tends to oscillate with the epoch numbers while the loss on the training set strictly decreases (Fig. 9a). However, the validation loss of ResNet-Patch/ResNet-NCRF has the similar tendency compared to its training loss (both of them are strictly decreasing).

**Fig. 9.**
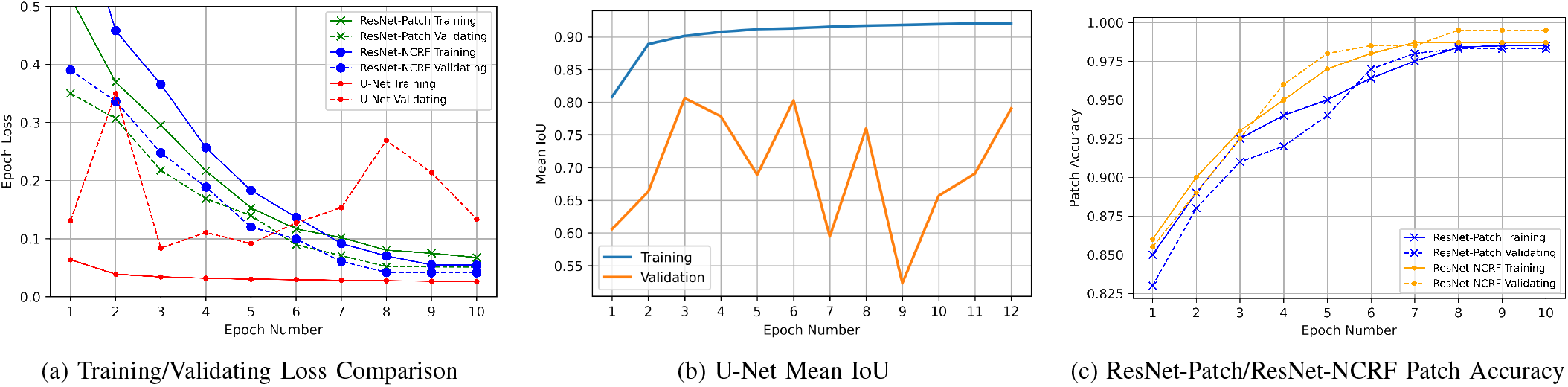
Training Process Comparison: we visualize the trends of training and validation to further compare ResNet-Patch/ResNet-NCRF with U-Net.

To further analyze the oscillation issue of U-Net and compare it with ResNet-Patch/ResNet-NCRF, we select mean IoU of training and validating as performance metric as U-Net is a pixel-level architecture that outputs pixel-wise masks for GM and WM. Fig. 9b shows its mean IoU results on the validation set tend to oscillate across the number of trained epochs while the results on the training set strictly increase (Fig. 9b). Besides, the mean IoU of training is over 0.9 the mean IoU for validation set is at most 0.8, even less than 0.55 at 9th epoch of training. Therefore, we performed five-fold cross-validation on the training and validation set to determine the optimal early stopping point for the number of training epochs. Then we trained the model for that number of epochs on the combined training-validation set and evaluated on the test set. The number of training epochs we obtained is 5 for U-Net.

On the other hand, ResNet-Patch/ResNet-NCRF is a patch-level classification problem, so we choose to examine the patch-level accuracy in the training/validating process. Fig. 9c shows that the classification accuracy of ResNet-Patch/ResNet-NCRF continues to increase for both training and validation sets. Both achieve 0.96 accuracy or above after 6 epochs, indicating that their results are very close without any oscillations or instability.

### B. Segmentation Visualization

Fig. 10 shows the segmentation visualization of FCN, U-Net, ResNet-Patch and ResNet-NCRF on hold-out AD and NAD cases separately. WSI-16 is a AD case while WSI-30 is a NAD case. In Fig. 10, our methods segment the whole WSIs into three areas: GM, WM, and background, which are indicated by cyan, yellow, and black, respectively. The mask of FCN (Fig. 10c, 10h) indicates FCN is not able to distinguish the tissue from background well (as shown in the top and bottom portions of the image where FCN detects GM when no tissue is present) while U-Net (Fig. 10b, 10g), ResNet-Patch (Fig. 10d, 10i), and ResNet-NCRF (Fig. 10e, 10j) can easily segment the tissue from background. The segmentation masks of U-Net ResNet-Patch, and ResNet-NCRF are visually the same as ground-truth annotations generated by trained personnel.

**Fig. 10.**
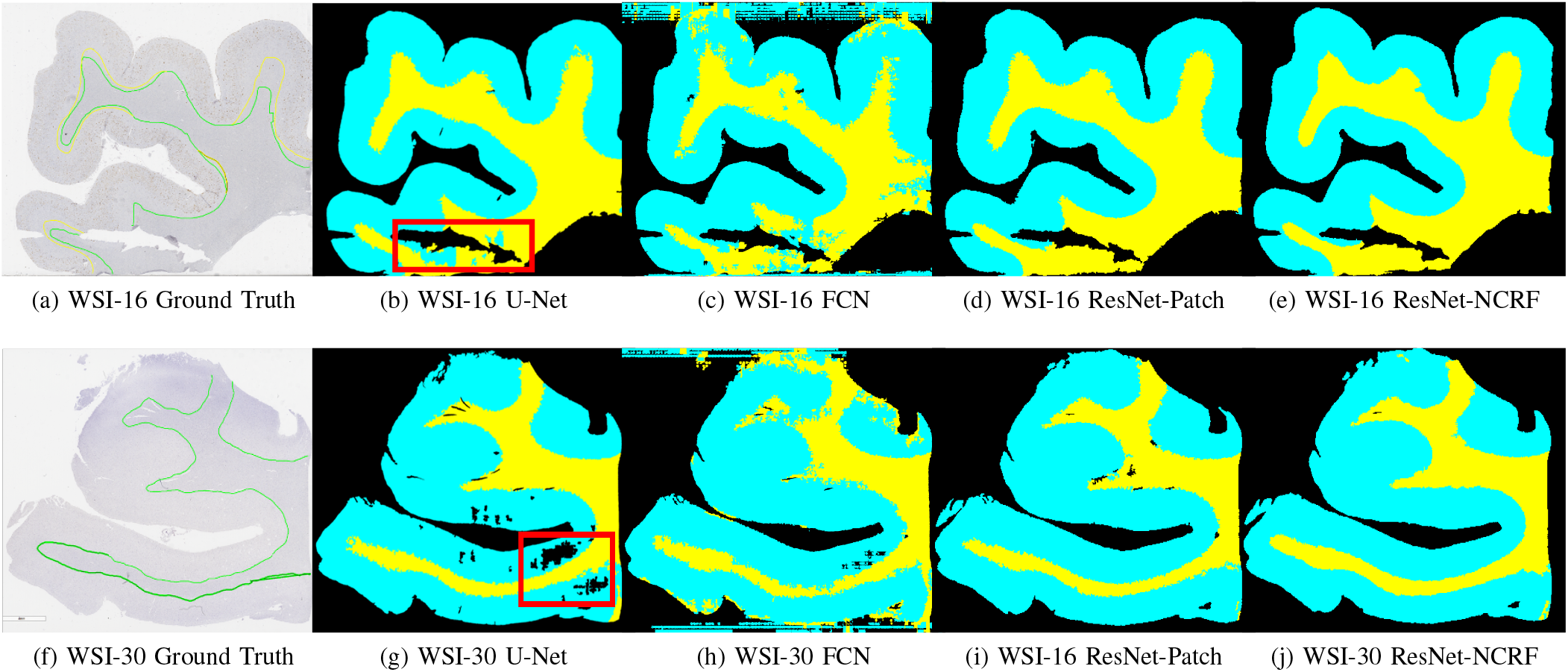
Segmentation masks visualization: WSI-16 is a AD case (top panels) and WSI-30 is a NAD case (bottom panels). GM, WM, and background are indicated by cyan, yellow, and black, respectively.

Examining the red block in Fig. 10b, some blue dots are inside of yellow area (WM), indicating some areas of WM are misclassified as GM by U-Net. To further evaluate this area, we provide the zoom-in details as shown in Fig. 11: these areas are from WM and they may be folds or torn off tissue, which misleads U-Net to a wrong prediction. But ResNet-Patch/ResNet-NCRF can predict correctly in these areas, which shows the robustness of them even in imperfect areas such as artifacts of tissue slides. The same in Fig. 10g, there are some black areas in GM, which means U-Net classifies these pixels as background rather than GM. However, in the same area of Fig. 10d, 10i, ResNet-Patch’s mask is closer to our ground truth, so is ResNet-NCRF (Fig. 10e, 10j). To compare the difference between the mask of ResNet-Patch and ResNet-NCRF, we find the boundary between GM and WM provided by ResNet-NCRF is smoother compared to that from ResNet-Patch, which verifies the function of NCRF layer to effectively model spatial information.

**Fig. 11.**
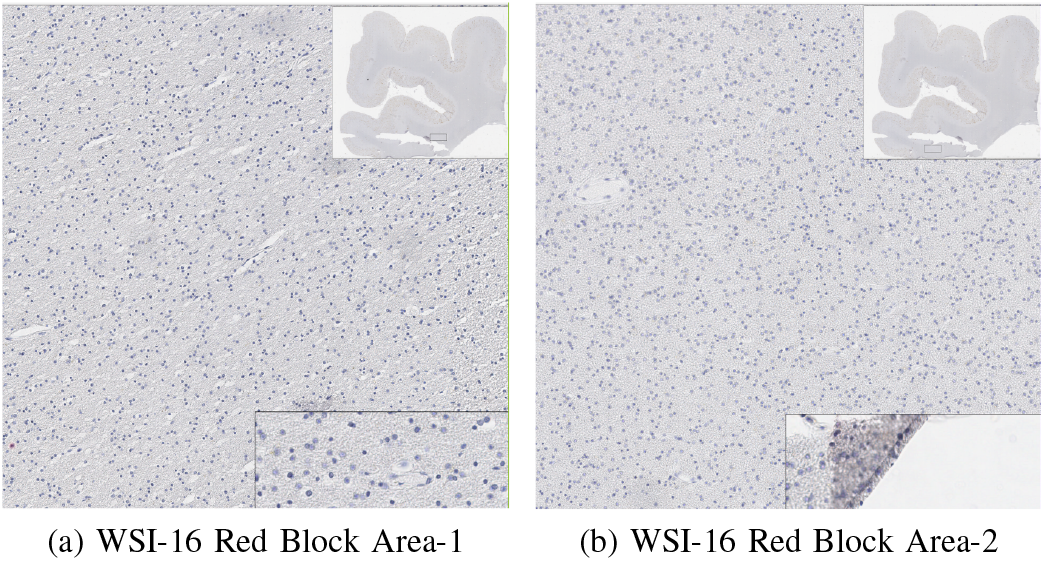
The zoom-in details of the areas within the red block in WSI-16.

### C. Grad-CAM Interpretation

#### 1) ResNet-18

To investigate the differential morphologies corresponding to GM and WM separately, we use Gradient-weighted Class Activation Mapping (Grad-CAM) [20] to generate a coarse localization map where the relevant regions for predicting the concept are highlighted. Note that both ReseNet-Patch and ResNet-NCRF has the same backbone, namely ResNet-18 as the feature extractor. Fig. 12 provides visual explanations of what features ResNet-18 are deemed important for differentiating GM and WM for both AD and NAD cases, respectively.

**Fig. 12.**
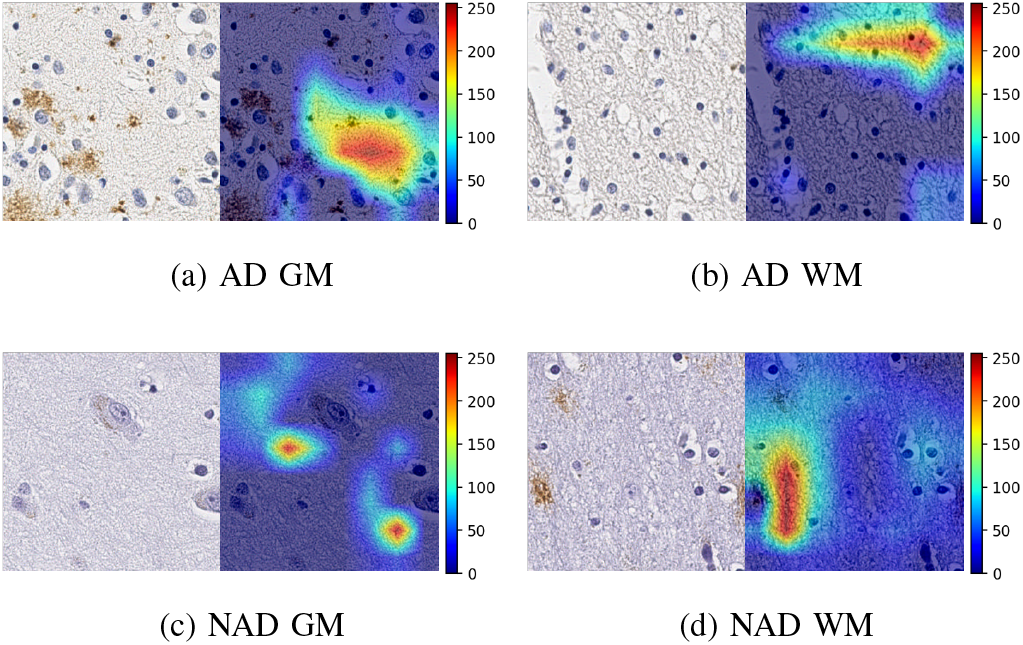
ResNet-18 Grad-CAM on selected patches from both AD and NAD cases.

From Fig. 12, we can see the energy around plaques (brown pixels in Fig. 12a) and cells is relatively lower, while the highest energy is focusing on the textures of brain tissue. Based on pathology differences between GM and WM, one hypothesize is the ResNet-18 gravitates towards selecting areas that are composed of more randomly associated fibers (i.e. neuropil consisting of dendrites in GM, and more organized myelinated axons in WM). This is interesting as GM is also comprised of a more heterogeneous cell population (i.e. neurons and glial cells) while the predominate cell type in WM is oligodendrocytes. As a result, ResNet-18 not only seems to ignore pathological hallmarks of Alzheimer’s, such as Amyloid-*β* plaques, but also other cellular components like neuronal and glial cells. Other abnormalities such as perivascular spaces, or artifacts are also avoided. The results have the potential to prove that our CNN feature extractor is relying on the differential textures from GM and WM rather than pathologies to determine the category of the patch, additional works with greater diversity (including other areas and stains) are needed. Therefore, our feature extractor is robust to datasets from both AD cases and NAD cases, which is clinically reliable and reasonable.

#### 2) U-Net

Similarly, we also apply Grad-CAM to our U-Net model to investigate the segmentation process. We generally find the convolutional layers near the end of U-Net’s contracting path captures more comprehensible features while the convolutional layers in the expansive path successively combine features from the contracting path and produce heatmaps that look more and more similar to the logits of the selected class. This Grad-CAM finding is supported by similar results of U-Net semantic segmentation on Cityscapes datasets [61]. In Fig. 13, we only show the Grad-CAM heatmaps of the 5th and 6th convolutional layers which are on U-Net’s third contracting level.

**Fig. 13.**
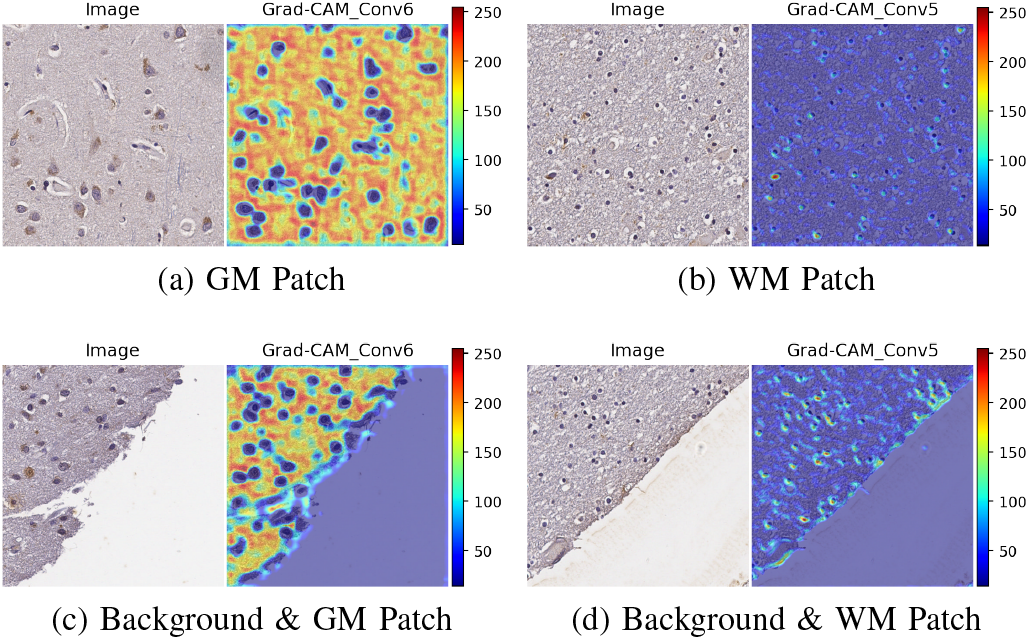
U-Net Grad-CAM results on selected convolutional layers.

As shown in Fig. 13, the 6th convolutional layer gives highest weights on tissue textures and near-zero weights on cell and neuron locations for patches containing GM (Fig. 13a,13c). For patches containing WM (Fig. 13b, 13d), the 5th convolutional layer gives highest weights on cell and neuron locations and low weights at other locations. These heatmaps indicate U-Net relies heavily on tissue textures when segmenting GM and on cell and neuron locations when segmenting WM.

## VI. Discussion and Conclusion

In this paper, we propose an entire pipeline automating the segmentation of GM and WM in WSIs of ultra-high resolution. This pipeline consists of two components: interpretable CNN module for segmenting GM/WM regions and post-processing module for reducing artifacts and residues existing in WSIs as well as generate XML annotations that are helpful for neuropathologists. The final output of XML annotations can be displayed on the original-size WSIs for neuropathological studies.

For the CNN module, we first investigate the application of U-Net and FCN to this medical image segmentation problem, then propose a novel segmentation mechanism tailored to solve the issue of ultra-high resolution. Our proposed model, ResNet-Patch/ResNet-NCRF, provides a memory-efficient mechanism to GM and WM with stable training convergence. We employ Grad-CAM to illustrate the interpretability for both U-Net and ResNet-Patch/ResNet-NCRF: the heat maps provide reasonable explanations on what CNN models rely on to distinguish GM from WM, which include cell textures and distinctive characteristics of GM and WM from a clinical perspective. Both U-Net and ResNet-Patch/ResNet-NCRF have the potential to provide more objective and cost-effective boundaries of GM/WM compared to manual segmentation. While U-Net is more time-efficient, ResNet-Patch/ResNet-NCRF is more memory-efficient with around 1% higher IoU results and more consistent performance among different brain tissue slides. Furthermore, ResNet-Patch/ResNet-NCRF is easy to train as U-Net has oscillates and instability during the training process. For the post-processing module, we demonstrate its effectiveness on removing the majority of residues and artifacts.

Although our sample size is small and based on only 1 anatomic area (temporal cortex), this automatic segmentation is a proof of concept that demonstrates accurate classification of WM despite the smaller ratio of WM region in the WSI. ResNet-Patch/ResNet-NCRF achieve similar results on both AD and NAD cases, which indicates our method is robust across multiple diagnostic groups (across AD and NAD cases).

A future direction is to evaluate our pipelines against a larger WSI dataset with various stains (such as hematoxylin and eosin stain), and brain regions differentially affected across the AD spectrum (hippocampus, frontal cortex, and visual cortex). We also plan to study and understand the further generalizability of this method by examining other disease entities such as Lewy body and vascular dementias.

